# 4-aminopyridine promotes accelerated skin wound healing via neurogenic-mediator expression

**DOI:** 10.1101/2021.10.31.465317

**Authors:** Mashanipalya G. Jagadeeshaprasad, Prem Kumar Govindappa, Amanda M. Nelson, Mark D. Noble, John C. Elfar

## Abstract

We discovered that 4-aminopyridine (4-AP), a potassium channel blocker approved by the FDA for improving walking ability in multiple sclerosis, greatly enhances skin wound healing. Benefits included faster wound closure, restoration of normal-appearing skin architecture, increased vascularization and reinnervation. Hair follicle neogenesis within the healed wounds was increased, both histologically and by analysis of K15 and K17 expression. 4-AP increased levels of vimentin (fibroblasts) and α-smooth muscle actin (α-SMA, collagen-producing myofibroblasts) in the healed dermis. 4-AP also increased neuronal regeneration with increased numbers of axons and S100^+^ Schwann cells (SCs), and increased expression of SRY-Box Transcription Factor 10 (SOX10). Treatment also increased levels of transforming growth factor-β (TGF-β), substance P and nerve growth factor (NGF), important promoters of wound healing. *In-vitro* studies demonstrated that 4-AP enhanced proliferation and migration of human keratinocytes and SCs, and that 4-AP enhanced cellular interactions between neuronal and non-neuronal cells to further accelerate wound healing. Thus, 4-AP enhanced many of the key attributes of successful wound healing and offers a promising new approach to enhance skin wound healing and tissue regeneration.

## Introduction

Enhancing the healing of skin wounds is an important and challenging medical problem, whether these are isolated lesions or components of larger traumatic injuries. As the major protective barrier between the sterile environment inside the body and the pathogen-rich external world, the evolutionary pressure to optimize efficient healing of skin wounds has been very high, and this process is normally effective. This long history of evolutionary selection for optimized skin wound healing raises the question of whether it is even possible to improve on normal healing processes. Moreover, if this is possible, can it be done with approaches that facilitate efficient movement from the laboratory to the clinic?

One of the challenges in enhancing skin wound healing is that many different cellular processes must work in concert for effective and comprehensive repair. Along with wound closure, proliferation, migration and/or differentiation of keratinocytes, fibroblasts, Schwann cells (SCs) and neurons, re-vascularization, regeneration of hair follicles and other cellular changes are all important in successful, healthy wound healing (e.g., (1–5)). In addition, the cells of the skin need to produce and establish a healthy microenvironment comprised of extracellular matrix and specific neuropeptides and growth factors, such as transforming growth factor-β (TGF-β) and nerve growth factor (NGF) (e.g., (1, 6–12)), all of which are necessary to promote wound healing.

Surprisingly, several calcium channel inhibitors, including amlodipine, verapamil, diltiazem, and nifedipine, and azelnidipine (13–16), promote some aspects of wound healing, raising the possibility that ion channel modulators may be useful in enhancing repair. Topical verapamil application increased the rate of wound closure, the density of fibroblasts and collagen bundles and the volume densities of blood vessels (14). Nifedipine and amlodipine enhanced skin tensile strength and amlodipine caused faster wound closure (15). Thus, these compounds enhance some aspects of normal wound healing, although not all components of the healing response were investigated in these reports.

A different class of ion channel inhibitor of potential interest for skin wound healing, due to its pro-reparative effects on Schwann cells and peripheral neurons, is the potassium channel blocker 4-aminopyridine (4-AP). It has been recently shown, by genetic manipulations, that normal function of Schwann cells is important for enabling effective skin wound repair (1). These outcomes raise the question of whether a pharmacological approach to enhancing Schwann cell recovery might have similar benefits. 4-AP is an interesting candidate for such studies, due to its ability to enhance recovery of both Schwann cells and neurons after peripheral nerve crush injuries (17–22). What 4-AP would do in skin injuries, however, is unknown and not predictable. 4-AP is best studied as an inhibitor of multiple voltage-gated potassium channels (23–27), which do not appear to have been studied in the context of skin wound healing. 4-AP also affects calcium levels, both by increasing intracellular calcium levels and activating high-voltage activated calcium channels (28–31). Therefore, 4-AP would be predicted to change calcium levels in ways that are opposite than occurs with calcium channel blockers, which can inhibit calcium entry post-injury, leading to the possibility 4-AP treatment would actually inhibit wound healing.

We now report that, in mice with full-thickness dorsal skin wounds, systemic 4-AP treatment caused more rapid wound closure, restoration of normal epidermal thickness, tissue structure, collagen levels, and increased vascularization and cell proliferation. Thus, 4-AP treatment enhances many of the key attributes of successful wound healing. These findings provide strong support for the hypothesis that 4-AP treatment can enhance both tissue repair and regeneration in acute injuries. The extensive prior studies on 4-AP safety and dosing (e.g., (17–20, 32)), and its approval for the treatment of multiple sclerosis (33–35), make this compound of great interest for rapid transition to clinical studies.

## Results

### 4-aminopyridine (4-AP) accelerates wound closure and enhances skin regeneration

Systemic treatment with 4-AP accelerated full-thickness skin wound closure. We created 5-mm diameter full thickness dorsal excisional wounds in 10-wk-old male C57BL/6 mice (36); mice were then randomized and treated with either saline or systemic 4-AP (33, 37–39) daily for 14 days (Figure 1A). Wounds were splinted with silicone rings to prevent wound contraction and were monitored by digital imaging for morphometry, percentage of wound healing and tissue regeneration on days 3, 5, 7, 9, 12 and 14 post wound (PWD) (Figure 1A).

**Figure 1.**
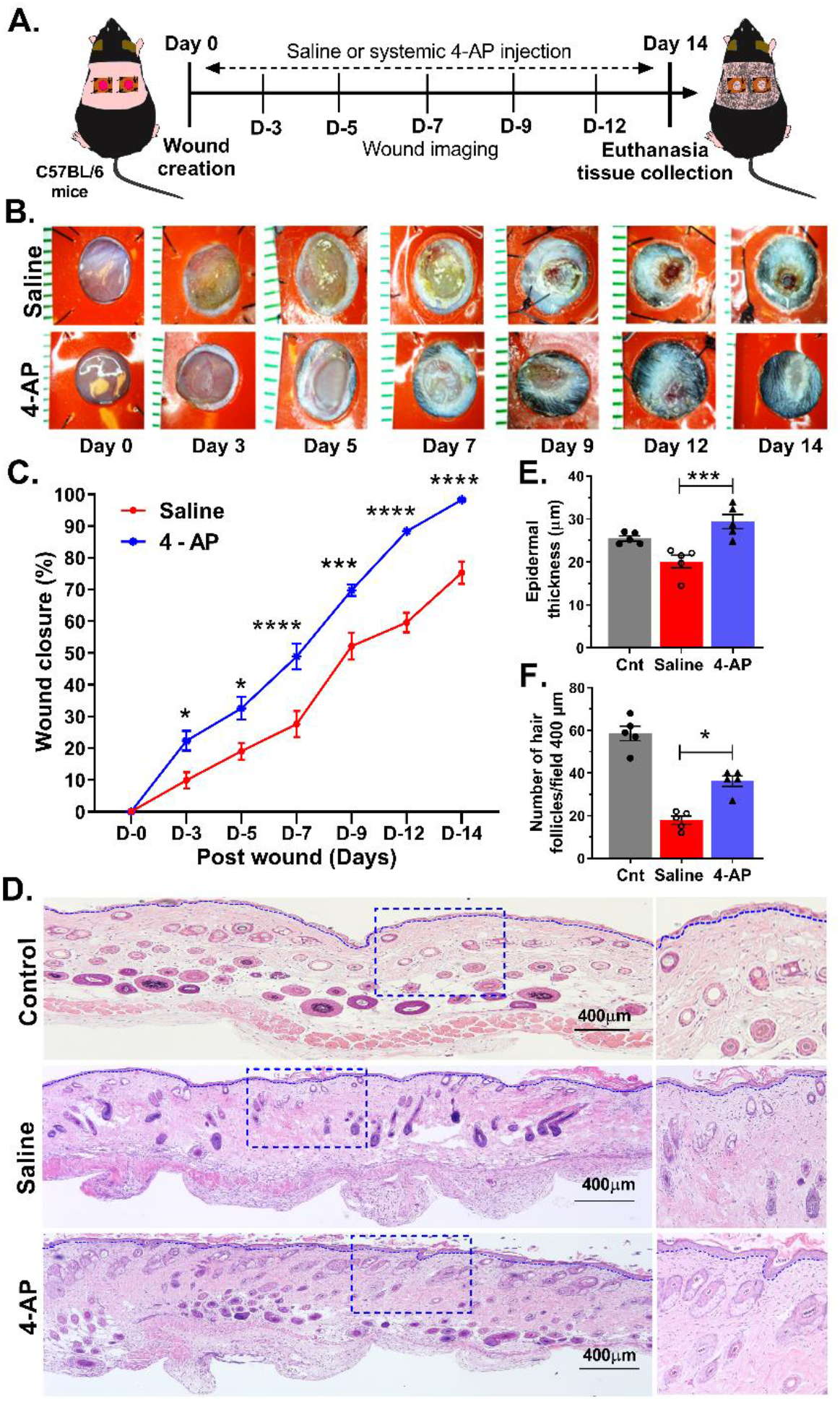
4-aminopyridine (4-AP) expedites wound closure and enhances skin regeneration. **(A)** Schematic illustration of experimental design to test the beneficial therapeutic effect of 4-AP in the C57BL/6 mouse splinted wound model. **(B)** Representative images of wound healing in control (saline-treated) and 4-AP treated mice at 0, 3, 5, 7, 9, 12, and 14 days post wounding (PWD). Scale bar, 1 mm. **(C)** Percent wound healing at each time point relative to the initial wound area in control and 4-AP treated mice. Value represents mean ± SEM, *n = 5* animals per group, with 2 wounds per animal, statistical significance indicated by asterisks (* = *P* between 0.01 and 0.05, *** = *P* between 0.001 and 0.0002, and **** = *P* between 0.0002 and 0.0001 vs. saline group), comparisons using two-way ANOVA (Sidak’s multiple comparisons test). **(D)** Representative images of H&E stained sections of normal control skin and full-thickness excisional wounds of saline-control and 4-AP treated skin tissue at PWD14. Scale bars = 400 µm. **(E)** Quantification of epidermal thickness in H&E stained sections by ImageJ software. **(F)** Quantification of the number of *de-novo* hair follicles within healed wounds Each image represents 5 images from 5 different mouse wounds and data are represented as mean ± SEM, *n = 5* animals per group, statistical significance indicated by asterisks (* = *P* between 0.01 and 0.05, and *** = *P* between 0.001 and 0.0002, vs. saline group).

We found that the extent of wound closure in 4-AP-treated mice was more than twice that of saline treated mice at PWD3 (Figure 1, B and C; p < 0.0199). Significant differences in wound closure were noted at every time point examined, including PWD14 (p < 0.0001). At this point, 4-AP-treated mice had complete wound closure, while open wounds were still present in saline-treated mice (Figure 1, B and C; p < 0.0001). Saline-treated mice had complete wound closure at PWD18, four days after complete wound closure in 4-AP mice (data not shown).

Histomorphometry analysis on PWD14 skin sections revealed that 4-AP treatment increased epidermal thickness compared with saline-treated mice, with the resulting epidermal thickness in 4-AP treated mice being restored to that of uninjured skin (Figure 1, D and E; p < 0.001) (1, 36, 40).

4-AP treated mice also showed a significant increase in wound-induced hair neogenesis (WIHN) when compared with saline-treated mice (Figure 1, D and F), a feature of successful skin regeneration (e.g., (41, 42)). 4-AP-treated mice exhibited a 1.8-fold increase in number of hair follicles compared to saline-treated mice (Figure 1F; p < 0.01).

### 4-AP increases keratinocyte number and epithelial stem-cell markers in healed wounds

Given the thickened epidermis in healed wounds with 4-AP (Figure 1, D and E), we investigated whether this was associated with increased numbers of keratinocytes and/or altered epidermal differentiation. We observed a 2-fold increase in the number of keratin-14 positive (K14^+^) keratinocytes in the epidermis and in the *de novo* hair follicles of 4-AP treated mice (Figure 2A) compared with saline-treated mice (Figure 2, A-C; p < 0.01). In contrast, expression of K10, a marker of epidermal differentiation, was not impacted by 4-AP treatment at this time point (Supplemental Figure 1, A and B) (43).

**Figure 2.**
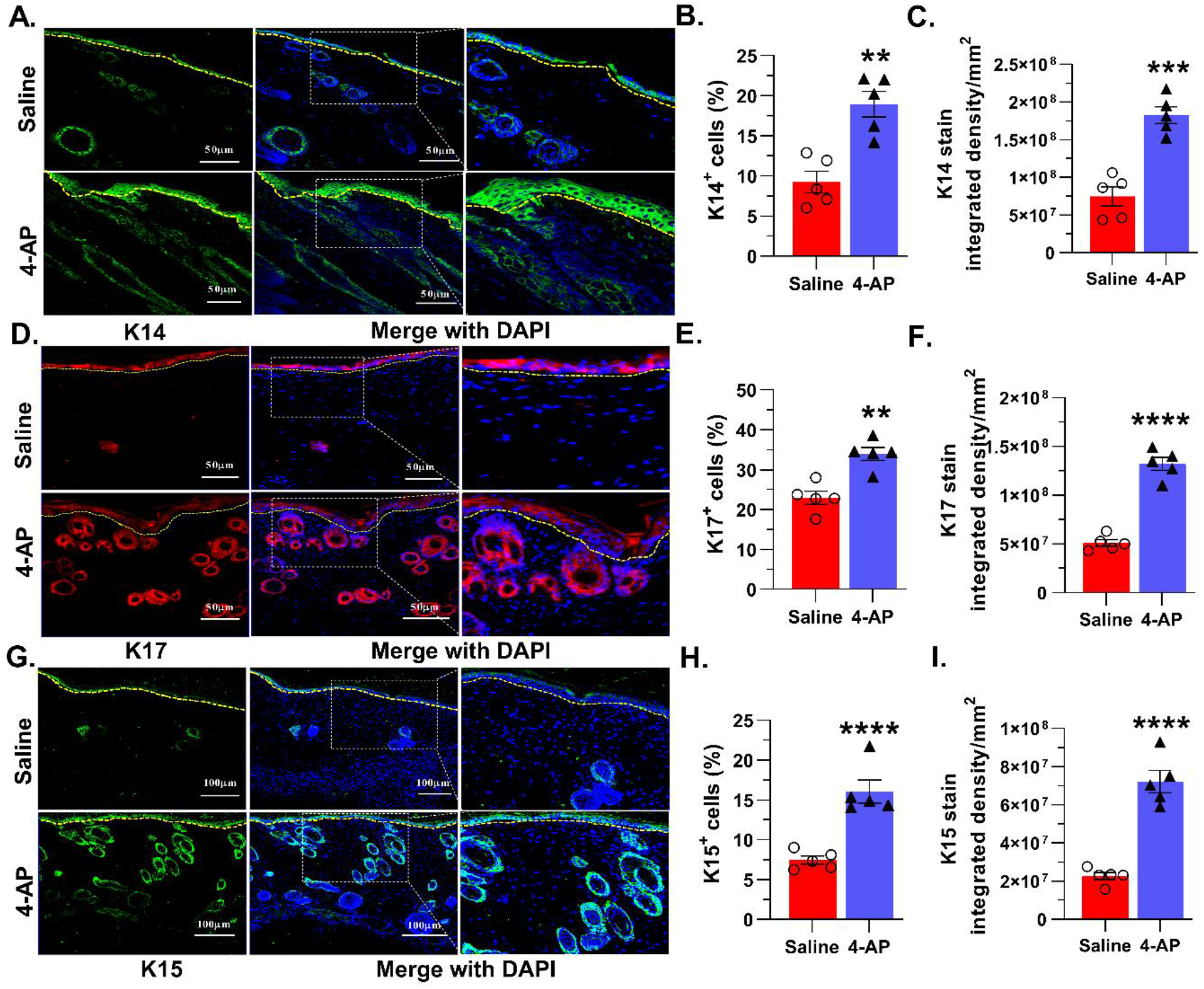
4-AP increases keratinocyte number and epithelial stem-cell markers in healing wounds. **(A)** Representative images of Keratin 14 protein expression by immunofluorescence in healed epidermis. K14 (green); DAPI (blue) denotes nucleus and dashed line denotes epidermal/dermal border. **(B and C)** Percent of K14^+^ cells and K14 protein integrated density in saline-control and 4-AP treated skin wounds at day 14. Scale bars = 50 μm. Data represents 20 images from 5 different mouse wounds and are shown as mean ± SEM, *n = 5* animals per group, statistical significance indicated by asterisks (** = *P* between 0.01 and 0.001, and *** = *P* between 0.001 and 0.0002 vs. saline). **(D)** Representative images of keratin 17 protein expression by immunofluorescence in saline and 4-AP treated skin wounds. K17 (red); DAPI (blue) denotes nucleus and dashed line denotes epidermal/dermal border. **(E and F)** Percent of K17^+^ cells and K17 protein integrated density in control and 4-AP treated skin wounds at day 14; Scale bars = 50 μm. Data represents 20 images from 5 different mouse wounds and are shown as mean ± SEM, *n = 5* animals per group, statistical significance indicated by asterisks (** = *P* between 0.01 and 0.001, and **** = *P* between 0.0002 and 0.0001 vs. saline). **(G)** Representative images of keratin 17 protein expression by immunofluorescence in saline and 4-AP treated skin wounds. K15 (green); DAPI (blue) denotes nucleus and dashed line denotes epidermal/dermal border. **(H and I)** Percent of K15^+^ cells and K15 protein integrated density in saline control and 4-AP treated skin wounds at day 14. Scale bars =100 μm. Data represents 20 images from 5 different mouse wounds and are shown as mean ± SEM, *n = 5* animals per group, statistical significance indicated by asterisks (****= *P* between 0.0002 and 0.0001 vs. saline).

In agreement with the increased numbers of hair follicles noted by histology (Figure 1, D and F), 4-AP increased the number of K17^+^ and K15^+^ cells and overall expression of these proteins further corroborating the increase in hair follicle number (41, 42, 44, 45). We observed a 1.5-fold increase in the percentage of K17^+^ cells (p < 0.001), and in K17 protein expression (Figure 2, D-F; p < 0.0001). Similarly, K15^+^ cells (p = 0.0005) and protein expression were increased with 4-AP treatment (Figure 2, G-I; p < 0.0001).

K14 and K17 expression also increased in the overlying epidermis in both saline and 4-AP-treated mice compared to uninjured skin, which is consistent with their known upregulation in response to wounding (Figure 2, A and D) (46).

### 4-AP treatment promotes increases in fibroblasts, myofibroblasts and transforming growth factor-β (TGF-β)

Fibroblast migration and maturation contribute to contraction, granulation, and proliferation phases of wound healing. A key marker of fibroblast differentiation is α-smooth muscle actin (α-SMA) which signifies fibroblast differentiation into collagen-producing myofibroblasts (e.g., (1, 3, 40)). We therefore investigated the effects of 4-AP treatment on fibroblasts and on expression of a known regulator of fibroblast differentiation, TGF-β.

To test whether 4-AP treatment altered fibroblast maturation during wound healing, we first performed Masson’s Trichrome staining to examine collagen deposition in the healing wound (Figure 3). This staining revealed elevated collagen deposition in 4-AP-treated mice compared to saline-treated mice (Figure 3, A and B; p = 0.0008), with collagen levels like those seen in normal tissue. This staining also revealed a tissue structure and collagen deposition pattern very much like that seen in normal tissue.

**Figure 3.**
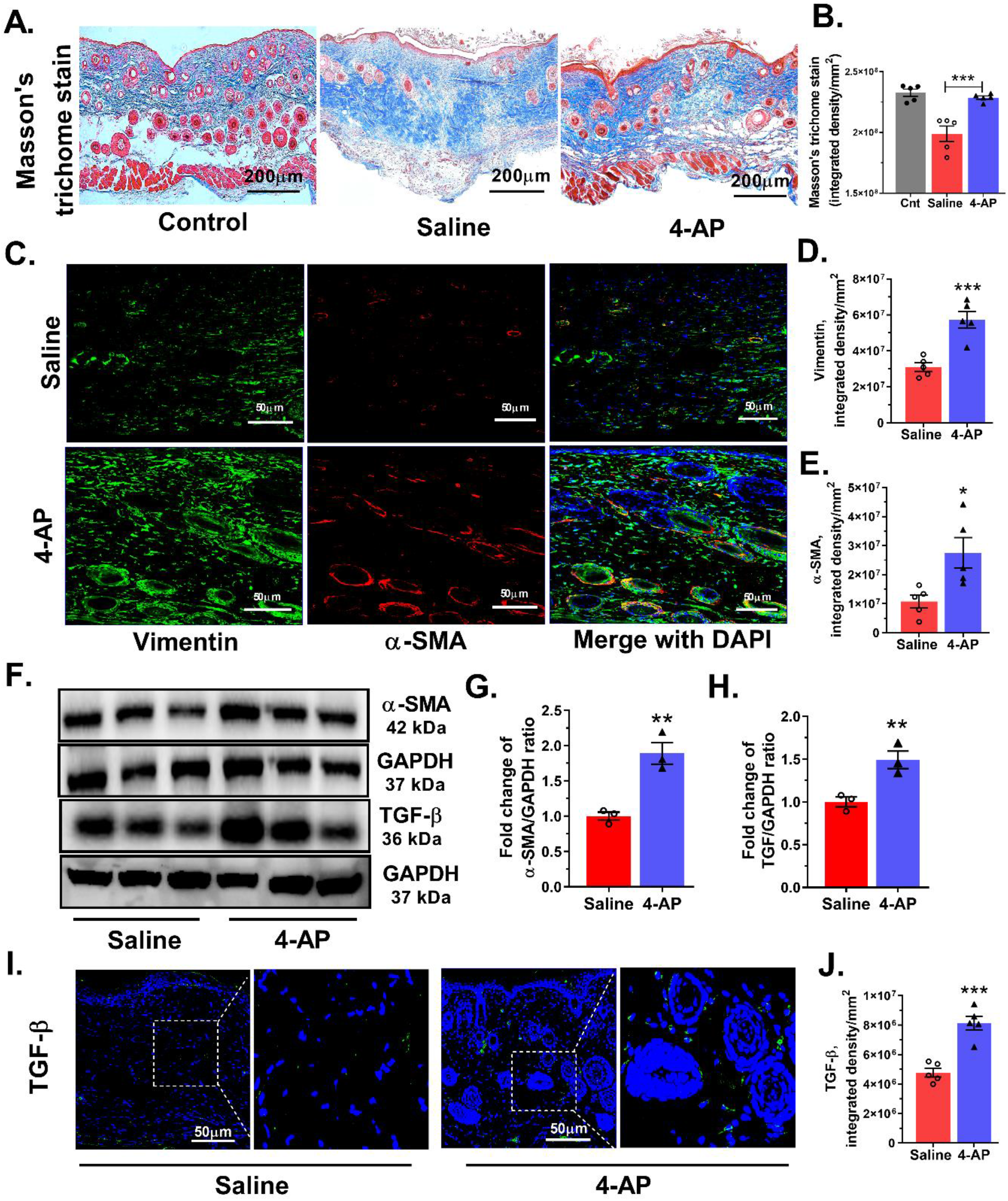
4-AP treatment increases fibroblast and myofibroblast numbers and levels of transforming growth factor-β. **(A)** Representative Masson’s trichrome stained images of healing control skin and full-thickness excisional wound of saline-control and 4-AP treated skin tissue at day 14 (PWD14). Scale bars = 200 μm. **(B)** Collagen density quantified as average blue pixel density per area in wound healing tissue harvested on day 14 (PWD 14). Data represents 10 images from 5 different mouse wounds and are shown as mean ± SEM, *n = 5* animals per group, statistical significance indicated by asterisks (*** = *P* between 0.001 and 0.0002vs. saline). **(C)** Co-immunofluorescence staining of vimentin (green), α-SMA (red) and nuclear stain (DAPI-blue) in saline control and 4-AP treated skin wound sections at day 14. Scale bars = 50 μm. **(D and E)** Quantification of vimentin and α-SMA protein staining intensity (integrated density). Data represent 20 images from 5 different mouse wounds and are shown as mean ± SEM, *n = 5* animals per group, statistical significance indicated by asterisks (* = *P* between 0.01 and 0.05, and *** = *P* between 0.001 and 0.0002 vs. saline). **(F)** Representative western blots of α-SMA and TGF-β levels. **(G and H)** Quantitation of α-SMA and TGF-β levels to normalized to GAPDH (fold change mean ± SEM, *n = 3* animals per group, statistical significance indicated by asterisks (** = *P* between 0.01 and 0.001vs. saline). **(I and J)** Representative images of TGF-β protein expression by immunofluorescence in wound tissue at PWD14. TGF-β (green), nuclei stained with DAPI (blue). Scale bars = 50 μm and TGF-β protein quantitative integrated density analysis. Data represents 20 images from 5 different mouse wounds and are shown as mean ± SEM, *n = 5* animals per group, statistical significance indicated by asterisks (*** = *P* between 0.001 and 0.0002 vs. saline).

4-AP treatment also increased the expression of fibroblast proteins, vimentin and α-smooth muscle actin (α-SMA). Immunofluorescence analysis revealed more vimentin^+^ fibroblasts and elevated vimentin levels in wound tissue from 4-AP-treated mice than saline-treated mice (Figure 3, C and D), which was consistent with western blot (WB) analysis for α-SMA expression (Figure 3, E and F). We also observed increases in α-SMA, which signifies fibroblast differentiation into collagen-producing myofibroblasts (e.g., (1, 3)). Increases were also seen in α-SMA protein (Figure 3, C, E-G; p = 0.0191).

TGF-β plays an important role in promoting myofibroblast differentiation (e.g., (1, 3, 9–11)), and we found significant increases in TGF-β protein expression with 4-AP treatment compared to saline treatment (Figure 3, F, and H-J; p = 0.0001).

### 4-AP promotes reinnervation and neuropeptide expression

Normal skin wound healing is also associated with increases in cell division and increases in non-dividing neurons. 4-AP treatment caused increases in both of these measures.

Expression of the proliferation marker, Ki-67, was significantly increased in mice treated with 4-AP compared with saline treated controls. The proportion of Ki-67^+^ cells within hair follicles and epidermis was increased 2.0-fold (Figure 4, A and B; p = 0.049).

**Figure 4.**
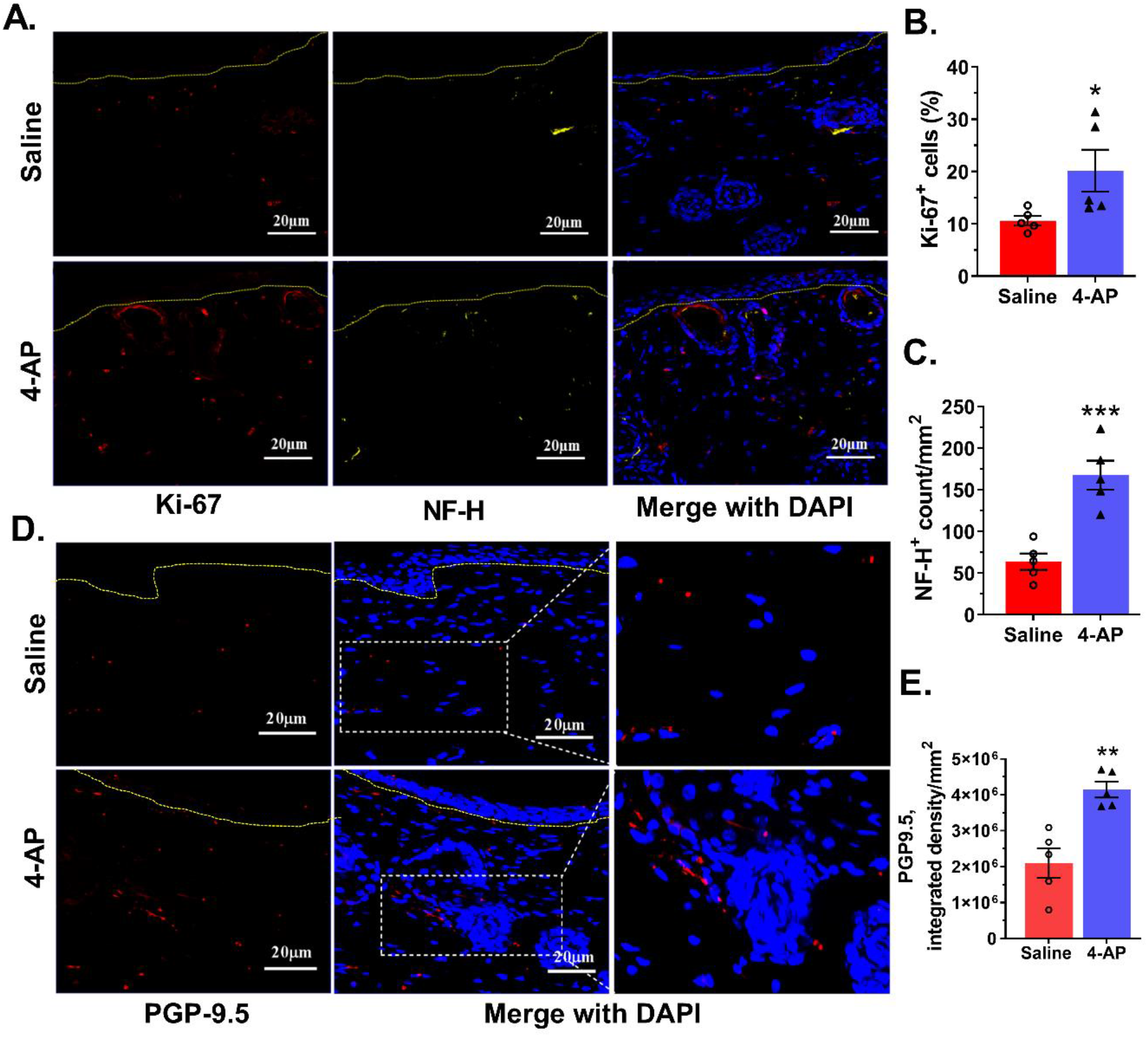
4-AP promotes division reinnervation and enhanced PGP-9.5 expression. **(A)** Representative images of co-immunostaining for Ki-67^+^ cells (red) and NF-H^+^ cells (yellow) surrounding *de novo* hair follicles in the healed wound at day 14 and dashed line denotes epidermal/dermal border. Scale bars =20 μm. **(B and C)** Ki67^+^ and NF-H^+^ cells were quantified. Data represents 20 images from 5 different mouse wounds and are shown as mean ± SEM, *n = 5* animals per group, statistical significance indicated by asterisks (* = *P* between 0.01 and 0.05, and *** = *P* between 0.001 and 0.0002 vs. saline). **(D)** Representative images of PGP-9.5 protein expression by immunofluorescence staining of healed wounds. PGP-9.5 (red) and nuclear stain DAPI (blue) and dashed line denotes epidermal/dermal border. Scale bars = 20 μm. **(E)** Quantification (integrated density) of PGP-9.5 protein expression. Data represents 20 images from 5 different mouse wounds and are shown as mean ± SEM, *n = 5* animals per group, statistical significance indicated by asterisks (** = *P* between 0.01 and 0.001 vs. saline).

The number of neurons in the skin of 4-AP-treated mice also was increased over that seen in saline-treated animals. Neuronal number was determined by staining with antibodies against high molecular weight neurofilament protein (NF-H) (e.g., (47, 48)). NF-H axonal counts were increased 2.5-fold in 4-AP-treated mice compared with saline-treated controls (Figure 4, A and C; p = 0.0008).

We also found that NF-H stained axons in the 4-AP treated mice were more often encountered in direct association with Ki-67^+^ hair follicles (Figure 4A) than in saline-treated controls, in agreement with observations that hair follicles are associated with sympathetically innervated *arrector-pili-muscles* (46). Thus, the association between nerve function and hair follicle stem cells under the influence of 4-AP supports reinnervation as a possible factor in the formation of *de-novo* hair follicles during wound healing.

Another example of the ability of 4-AP to restore aspects of skin structure like that seen in uninjured tissue was revealed by staining for protein gene product 9.5 (PGP-9.5), a neuronal peptide associated with wound healing (49). Fourteen days post wounding, PGP-9.5^+^ nerve fibres in the healed wounds were twice as abundant in 4-AP treated mice, as reflected by increased amounts of PGP-9.5, as compared with saline treated mice (Figure 4, D and E; p = 0.0023). The levels of PGP-9.5 in 4-AP-treated mice were not significantly different from seen in uninjured skin tissue (Supplemental Figure 1, C and D). This suggests that 4-AP significantly increased the expression of PGP-9.5 during wound healing and that 4-AP likely enhances skin reinnervation.

### 4-AP increases numbers of Schwann cells (SC) and expression of markers of an early differentiation state

Schwann cells (SC) are critical players in wound healing and are associated with axons around hair follicles in the wound bed. In the setting of injury, SCs de-differentiate to a non-myelinating state and begin to secrete neurotrophins like NGF, a state marked by expression of p75-NTR (50–52). We found that the number of SCs was significantly increased in the wounds of 4-AP-treated mice (Figure 5, A and B). Analysis of expression of S100, a pan-SC marker (e.g., (53)), identified SCs within both the hypodermis and dermis of the healed wounds (Figure 5A). The number of SCs was 3-fold greater in 4-AP treated mice than in saline treated controls (Figure 5, A and B; p < 0.01). SCs were preferentially located around nerve bundles, as predicted by the known affiliation of SCs with nerve cells. 4-AP treatment also increased the expression of p75-NTR (Figure 5, A, and C-F; p < 0.01), which is thought to be expressed in S100^+^ cells as a marker of de-differentiation (e.g.,(1, 50, 54)).

**Figure 5.**
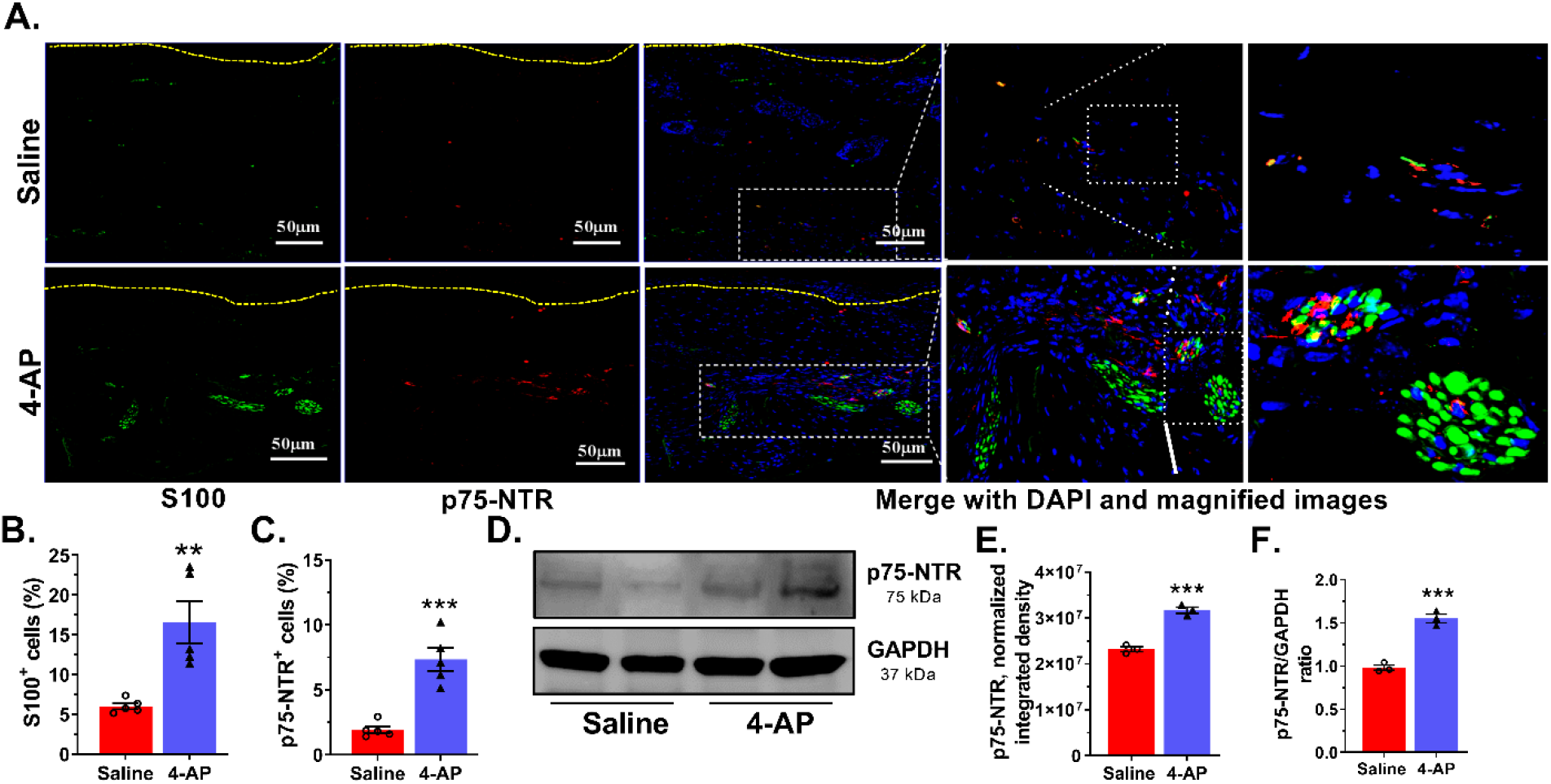
4-AP increases number of Schwann cells and expression of markers of an early SC differentiation state. **(A)** Representative images of co-immunostaining of S100 (green) and p75-NTR (red) in wound sections and dashed line denotes epidermal/dermal border. Scale bars = 50 μm. **(B and C)** Quantification of S100^+^ and p75-NTR^+^ expressing cells in healed wounds. Data represents 20 images from 5 different mouse wounds and are shown as mean ± SEM, *n = 5* animals per group, statistical significance indicated by asterisks (** = *P* between 0.01 and 0.001, and *** = *P* between 0.001 and 0.0002 vs. saline. **(D)** Representative image of western blot of p75-NTR and GAPDH. **(E and F)** Normalized integrated densities and ratio of p75-NTR protein expression represented as mean ± SEM, *n = 5* animals per group, statistical significance indicated by asterisks (*** = *P* between 0.001 and 0.0002 vs. saline).

We also found elevated expression of SOX10 and NGF in 4-AP treated mice. SOX10 is required for myelin production in SCs and elevated SOX10 expression promotes conversion of mesenchymal cells into p75-NTR expressing neural crest stem cells (NCSC) (e.g., (1, 55–57)). Conversely, depletion of SOX10 expression significantly delays wound healing and tissue regeneration (1). NGF plays a significant role in the wound healing process by inducing nerve sprouting from injured nerve endings (6–8, 58–60). NGF also promotes keratinocyte proliferation, and migration of dermal fibroblasts (7, 61). NGF also acts on non-neuronal cells to sensitize them to substance-P, which in turn further stimulates more NGF secretion, ensuring that keratinocytes can elaborate and respond to neuronal factors along with neurons (62, 63).

We found significantly increased SOX10 protein expression by both immunofluorescence and western blot analysis (Figure 6, A, B, E and F; p < 0.001). Substance-P and NGF protein expression was also increased in healed wounds from 4-AP treated mice compared to saline treated mice (Figure 6, A, C-G; p < 0.01). These factors, which are known to be associated with both nerve regeneration in the wound bed and accelerated healing (e.g., (1, 6–8, 58, 59, 63)), were increased within the wound with 4-AP treatment.

**Figure 6.**
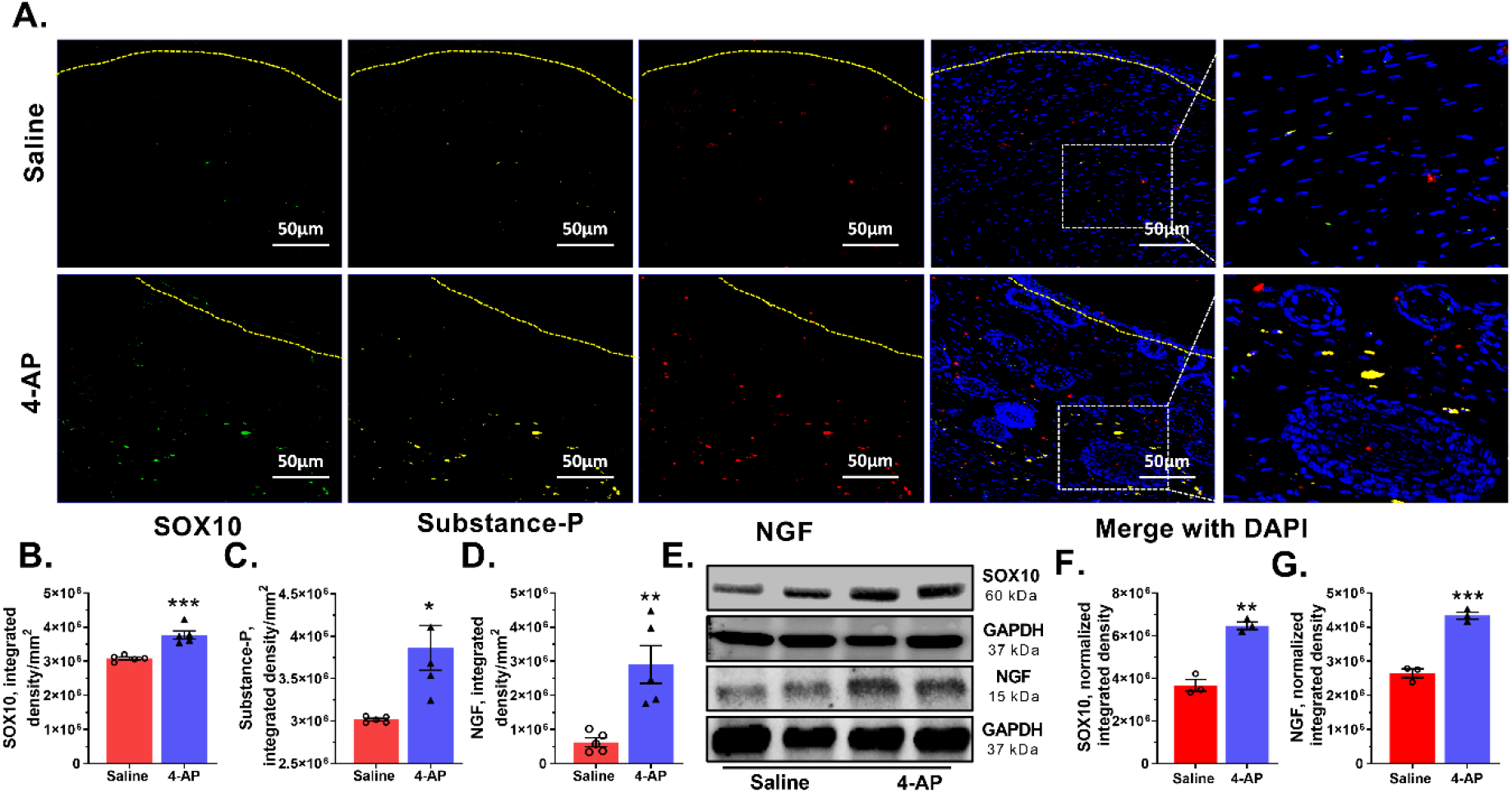
4-AP enhanced expression of transcription factors, neurotrophic factors, and neuropeptides associated with reinnervation. **(A)** Representative images of triple co-immunostaining of wound skin for the transcription factor SOX10 (green), neuropeptide substance-P (SP — yellow), nerve growth factor (NGF — red), and nuclear stain DAPI (blue) and dashed line denotes epidermal/dermal border. Scale bars = 50 μm. **(B-D)** Quantification of SOX10, substance-P and NGF expressing cells in healed wounds. Data represents 20 images from 5 different mouse wounds and are shown as mean ± SEM, *n = 5* animals per group, statistical significance indicated by asterisks (* = *P* between 0.01 and 0.05, ** = *P* between 0.01 and 0.001, and *** = *P* between 0.001 and 0.0002 vs. saline). **(E)** Representative image of western blot of SOX10, NGF and GAPDH. **(F and G)** Normalized integrated densities for SOX10, and NGF immunoblot. Mean ± SEM, *n = 3* animals per group, statistical significance indicated by asterisks (** = *P* between 0.01 and 0.001, and *** = *P* between 0.001 and 0.0002 vs. saline).

### 4-AP promotes neo-angiogenesis in granulation tissue

Neo-angiogenesis is necessary to provide nutrients and oxygen to healing wounds. To assess whether 4-AP treatment enhanced wound neo-vascularization, we performed immunofluorescence staining for the endothelial specific marker, CD31 (e.g., (40, 64)).

We observed larger and more abundant blood vessel networks in dermal tissue from 4-AP treated mice than saline treated mice (Supplemental Figure 1E). Although the number of blood vessels in 4-AP-treated mice was still less than in uninjured tissue, vessels in the newly healed tissue were notably larger. Quantification of CD31 intensity revealed statistically significant increases in blood vessels with 4-AP treatment compared to saline treatment (Supplemental Figure 1F). These results suggest that 4-AP treatment enhances neo-angiogenesis – a likely contributing factor in improved wound healing and tissue regeneration.

### 4-AP effectively stimulates proliferation and migration in primary cultures of human skin derived primary cells in-vitro

We next found that effects of 4-AP on keratinocytes and SCs *in-vitro* were similar to outcomes observed *in-vivo*, suggesting 4-AP may act directly on these cell types. In these experiments, we cultured primary, normal human epidermal keratinocytes (NHEKs) and dermal SCs in the presence or absence of 4-AP (65–67).

Following initial isolation from human foreskin, we confirmed the purity and identity of keratinocytes, and dermal SCs by immunohistochemistry using characteristic protein markers (Supplemental Figure 2, A and B). There was a modest decrease in cell viability at higher 4-AP concentrations (>10mM), but no obvious impact on cell viability at 2 mM concentrations of 4-AP (Supplemental Figure 2, C and D) (47). To determine the effect of 4-AP on wound healing *in vitro,* automated wound scratch assays were performed (e.g., (47, 59, 68, 69)) on confluent monolayers of keratinocytes, and SCs, with and without 1 mM 4-AP, a standard 4-AP dose used in studies *in-vitro,* (e.g., (47, 68)). 4-AP exposure accelerated scratch closure and keratinocyte migration (Figure 7, A and B; p < 0.01; and Supplemental Movie 1-2) within 3 hours, with complete scratch closure occurring at 18 hours. In contrast, control cultures not exposed to 4-AP closed at 32 hours (Figure 7, A and B).

**Figure 7.**
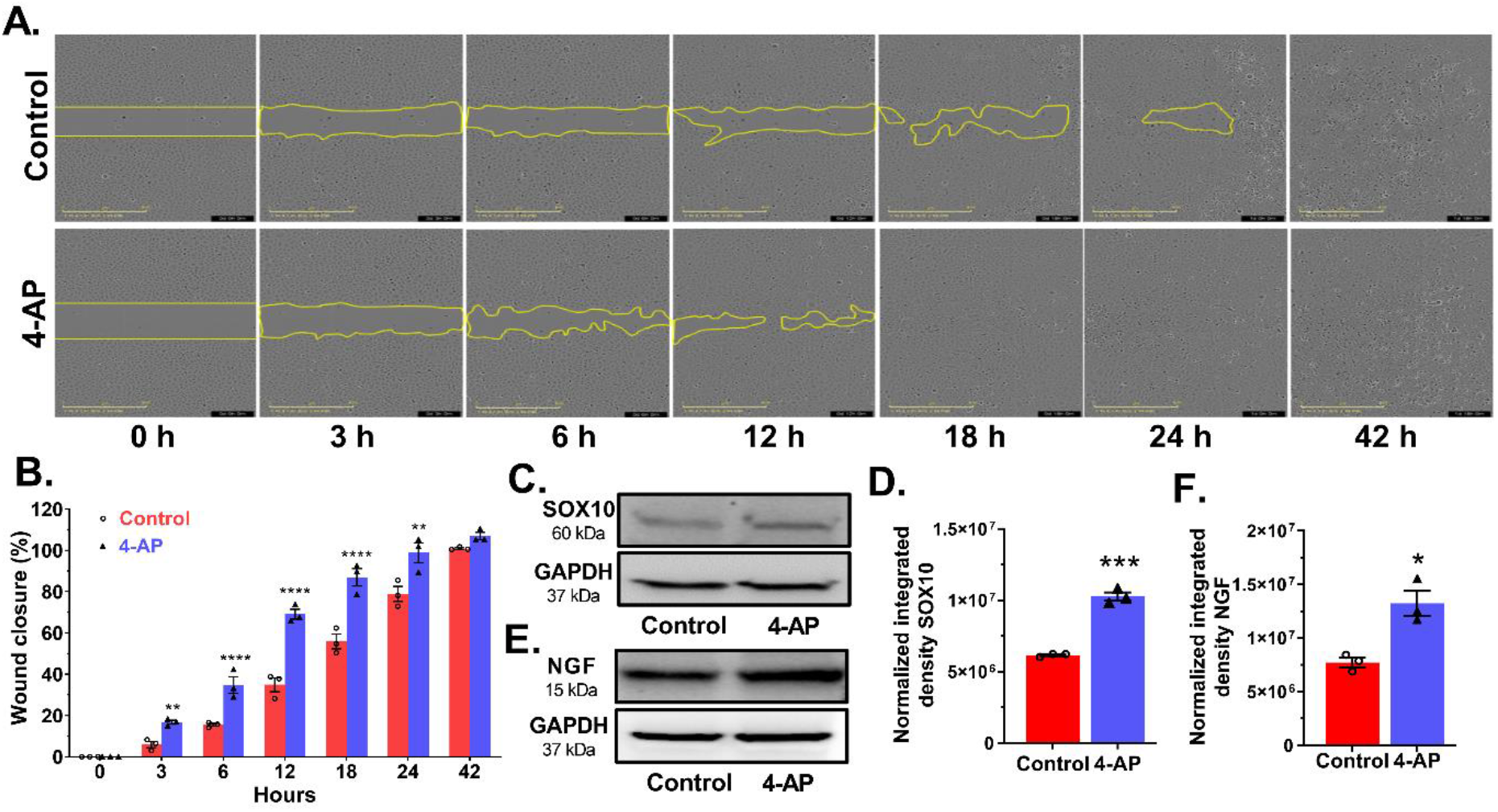
4-AP accelerates *in-vitro* keratinocyte wound closure. **(A)** Representative images of *in-vitro* keratinocyte scratch assays with 4-AP and vehicle control at indicated time points. The yellow lines indicate the wound borders at the beginning of the assay and were recorded every hour until 42 hours. Scale bar =100 µm. **(B)** The relative percentage of wound closure was calculated as the ratio of the remaining wound gap at the given time point compared to time 0. Each image represents 9 images from 3 biological replicates and data represented as mean ± SEM, *n= 3* biological replicates per group, statistical significance indicated by asterisks (** = *P* between 0.01 and 0.001, and **** = *P* between 0.0002 and 0.0001 vs. control). **(C -F)** A representative western blot and normalized integrated densities for SOX10, and NGF. Mean ± SEM, *n = 3* biological replicates per group, statistical significance indicated by asterisks (* = *P* between 0.01 and 0.05, and *** = *P* between 0.001 and 0.0002 vs. control).

SOX10 and NGF expression in 4-AP treated keratinocytes were both increased (Figure 7, C-F), suggesting that 4-AP promotes a more proliferative, stem-like phenotype in keratinocytes that contributes to accelerated scratch closure (55, 59). Similarly, in SCs, 4-AP also accelerated scratch closure, with 80% closure occurring before 11 hours with 4-AP treatment compared with 23 hours without treatment (Supplemental Figure 3, A and B; and Supplemental Movie 3-4).

### 4-AP treatment enhanced the co-cultured cell interaction and accelerated wound closure in-vitro

We next found that co-culturing cells improved scratch closure rates and that 4-AP further accelerated these effects. Given that keratinocytes, and SCs cells all interact during wound healing *in-vivo*, we conducted co-culture experiments with pairs of cell types with and without 4-AP treatment. Keratinocytes co-cultured with SCs in a ratio mimicking that of the epidermal skin (10:1 ratio of keratinocytes:SCs) closed a scratch within 15 hours with 4-AP treatment. This was significantly faster than closure without treatment, which took 20 hours. At 15 hours, the percentage of wound closure was at 98% in 4-AP treated co-cultures but only 80% in non-treated co-cultures (Figure 8, A and B; p < 0.0001; and Supplemental Movie 5-6).

**Figure 8.**
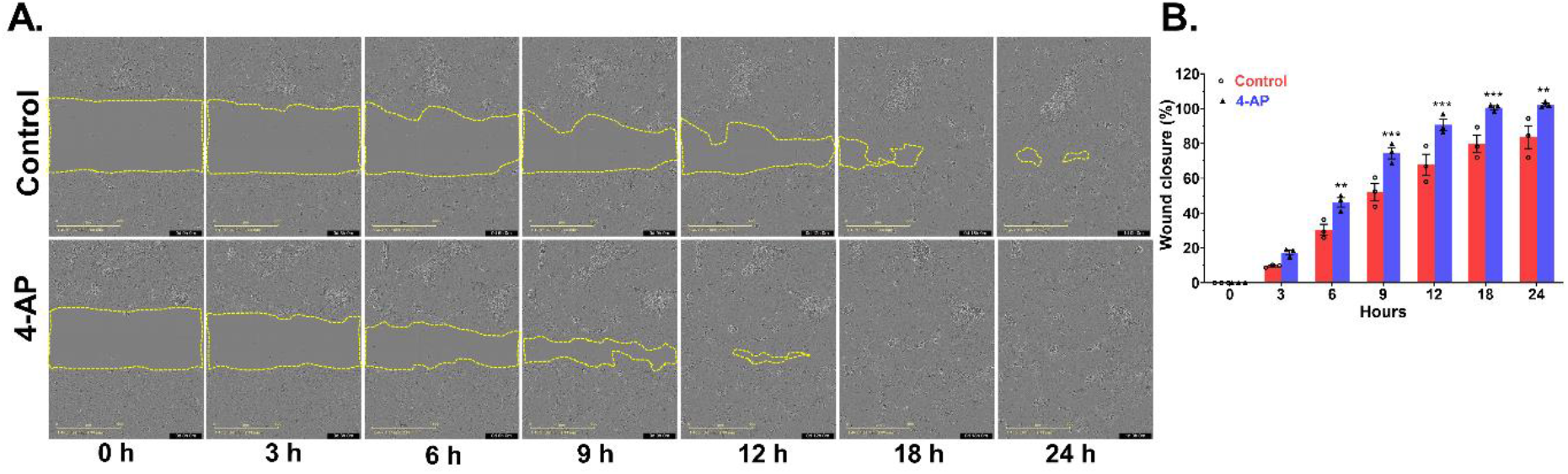
4-AP treatment enhanced the co-cultured cell interaction and accelerated wound closure and mechanistic insight of 4-AP effect on keratinocyte wound closure. **(A)** Representative images of *in-vitro* co-cultured keratinocytes and Schwann cells in scratch assays with 4-AP and vehicle control at indicated time points. The yellow lines indicate the wound borders at the beginning of the assay and were recorded every hour until 24 hours. Scale bar = 100 µm. **(B)** The relative percentage of wound closure was calculated as the ratio of the remaining wound gap at the given time point. Each image represents 9 images from 3 biological replicates and data represented as mean ± SEM, *n=3* biological replicates per group, statistical significance indicated by asterisks (** = *P* between 0.01 and 0.001, and *** = *P* between 0.001 and 0.0002 vs. control).

## Discussion

We found that, in mice with dorsal skin wounds, systemic 4-AP treatment caused more rapid wound closure, restoration of normal epidermal thickness, tissue structure, and collagen levels, and increased vascularization and cell proliferation. Hair follicle numbers were also increased in 4-AP treated mice, as determined histologically and by analysis of K15 and K17 expression, as were numbers of K14^+^ keratinocytes. Levels of vimentin (a marker of fibroblasts) and α-SMA (a marker of collagen-producing myofibroblasts) were increased, as were the numbers of α-SMA^+^ cells. 4-AP treatment also increased numbers of axons and S100^+^ Schwann cells, and increased expression of SOX10. Levels of several factors involved in skin repair also were increased (i.e., TGF-β, NGF and Substance P). Thus, 4-AP enhanced many of the key attributes of successful wound healing. As an already approved therapeutic agent, 4-AP appears to offer a promising new approach to wound healing and skin regeneration.

Enhancing normal skin wound healing is a difficult challenge because of how effective this process already is. Expediting and enhancing the wound healing process, however, could help in restoring the skin barrier even more quickly and in mitigating against infection complications. The many millions of years of evolutionary optimization of this process has resulted in a complex orchestration of division, migration and differentiation of multiple cell types, processes that depend on multiple growth factors and extracellular matrix components. Interference with any combination of these individual contributors can inhibit normal healing, emphasizing the extent to which healing requires the orchestrated interplay of many different components (including, for example, contributors as diverse as Wnt signaling, hair follicle development, β1 integrins, keratinocyte migration, extracellular matrix, macrophages, neovascularization, multiple growth factors, Schwann cells, appropriate TGF-β signaling, HIF-1 regulation, control of inflammation, etc. (e.g., (1, 2, 4, 9–11, 41, 70–87)). Despite the advances made in understanding the events required for effective skin wound healing, it has proven difficult to identify approaches that can enhance normal repair processes.

The ability to enhance normal wound healing with 4-AP is surprising for a number of reasons. 4-AP has the ability to enhance multiple processes associated with successful wound healing simultaneously. Our results are surprising because interfering with ion channel function seems less specific than more targeted approaches of cell transplantation or growth factor/cytokine manipulation during wound healing. Nonetheless, several calcium channel blockers also can enhance at least some aspects of skin would repair (13, 14, 16, 88). Only a subset of the outcomes in the present study were examined in these previous studies, which prevents a full comparison of potency of 4-AP with these other calcium channel modulators. Nonetheless, these earlier studies do support the idea that modulating function of at least some ion channels is a potentially useful approach to enhancing wound healing. That said, the benefits obtained with calcium channel blockers also raise concerns about whether 4-AP would be effective for this purpose, or whether it might even have adverse effects. This is because treatment with calcium channel blockers, such as verapamil, acts to inhibit calcium entry post-injury. In contrast, several studies on 4-AP predict it might increase intracellular calcium levels (e.g., (28, 30, 47)). Therefore, while it is intriguing that different classes of ion channel modulators can enhance healing of normal skin wounds, the prior indications that 4-AP and calcium channel blockers might have opposite effects on intracellular calcium levels makes it difficult to offer a unifying hypothesis for these observations. Regardless of whether or not a common mechanism exists for the effects of these two classes of ion channel modulators, the data from our studies and from previous studies on calcium channel blockers indicate that treatment with ion channel modulators offers a promising approach for enhancing repair of skin wounds.

The ability of 4-AP to enhance regenerative processes post-injury was discovered only recently, and the present work provides still stronger support for the potential utility of this compound as a pro-regenerative agent than our previous findings. In our previous studies on peripheral nerve crush injuries, we found that 4-AP caused a more rapid return of motor function, a more rapid and greater return of nerve conduction and increases in numbers of neurons and in myelination (17). The more rapid restoration of myelination and physiological function provide evidence for pro-regenerative effects of 4-AP treatment. Some of the benefits observed in our previous studies could have been due, however, to protective effects of this drug, as recently reported in models of CNS damage and in multiple sclerosis patients (47, 89, 90). However, in the case of full-excisional skin wound healing, outcomes such as those reported herein clearly require tissue regeneration.

The utilization of 4-AP to promote tissue repair is a qualitatively new use for this well-studied and clinically-utilized drug. The positive tissue regenerative effects of 4-AP in the skin are very novel from the extensively studied ability of this drug to provide transient symptomatic relief, without evidence of regenerative changes, in chronic neurological illnesses and injuries as multiple sclerosis, myasthenia gravis, cerebellar gait ataxias, downbeat and upbeat nystagmus and spinal cord injury (e.g., (91–106)). In neurological cases, the behavioral benefits of 4-AP are lost when treatment is terminated and the drug is cleared from the body. In contrast, the regenerative changes seen in acute traumatic injuries suggest that, in these settings, 4-AP treatment provides benefits that can endure long after treatment ends.

Whether any single target of 4-AP is most critical in skin wound healing, or whether modulation of multiple processes by this drug is critical, will be an interesting but challenging puzzle to solve. Wound healing processes are intricately connected and coordinated. We initiated the present experiments based on our previous findings that 4-AP treatment of acute peripheral nerve injuries enhanced SC and neuronal function post-injury and to the importance of SC and neuron function in wound healing (1, 6, 17, 18, 32, 107). The multiple effects of 4-AP treatment, however, suggest that SCs and neurons were not the only components of successful wound healing impacted by this drug. For example, NGF, Substance P and TGF-β levels were all elevated by 4-AP treatment, and each of these has been shown to have pro-reparative effects in skin wound lesions (e.g., (8–11, 108–113)). Our *in-vitro* studies also indicated that 4-AP can have direct effects on keratinocyte migration and expression of SOX10 and NGF by these cells, indicating that the actions of 4-AP are not restricted to neuronal cell types. Consideration of direct molecular targets of 4-AP is also complex, and includes multiple potassium and calcium channels and may also include the ability to sequester intracellular calcium (e.g., (114–116)). Skin cells, including keratinocytes, are dependent on Ca^+2^ signaling for proper differentiation programs further increasing the complexity of this problem. Elucidating the exact molecular mechanism(s) by which 4-AP enhances skin wound repair is challenging, but nonetheless, its translatability to the clinic is very promising. In contrast with the challenges in mechanistic analysis of the effects of 4-AP, the translation of our findings to clinical studies is relatively straightforward. 4-AP fits the very definition of drug repositioning: using existing clinically useful compounds in novel applications. The ability to accelerate skin wound healing and skin regeneration is novel. Extensive prior studies on 4-AP safety and dosing (e.g., (24, 26, 92, 93, 99, 117)), and its FDA approval in 2010 for the treatment of multiple sclerosis, make the transition of this compound from laboratory back to the clinical arena for wound healing therapy relatively straightforward. Indeed, our findings that 4-AP treatment can be used to distinguish between incomplete and complete peripheral nerve injuries have already transitioned to clinical trials on the diagnosis of such injuries (https://clinicaltrials.gov/ct2/show/NCT04026568)(17, 20). Moreover, our findings that 4-AP treatment enhances functional recovery from peripheral nerve damage have transitioned to clinical trials focused on enhancing recovery from peripheral nerve damage associated with radical prostatectomy, as manifested by urinary incontinence and erectile dysfunction (https://clinicaltrials.gov/ct2/show/NCT03701581)(17, 20, 32). Thus, the possibility of bringing the present findings forward for clinical examination seems a promising one. In a broader sense, the large numbers of available drugs that modify ion channel function (e.g., (118–120)) offer multiple additional candidates of interest their potential use in regenerative medicine.

## Methods

### Study design

The primary objective of this study was to investigate the possible therapeutic effect of 4-AP in enhancing skin wound healing and tissue regeneration in C57BL/6 male mice. Mice were age-matched and randomized to treatment groups: systemic 4-AP or saline. The number of animals (n) needed was calculated based on the conservative use of animals for the least sensitive data type and was determined based on a desired power level greater than 80% and a required P<0.05. The total we started with was 8 animals per group. The probability of mice dying in unrelated experiments was 8%, and was also an 8% chance of the splint coming out (nonadherent) from the wounded mice, and we excluded such mice from the study. The selected number of animals (n=5; 2 wound/animal; and total 10 wounds) used per group qualifies for the classical pre-hoc power analyses. Data were generated by microscopic analysis of immunohistochemistry, immunofluorescence on fixed skin sections, and immunoblotting of tissue and cell lysates. Five animals with 10 wounds were used in functional wound closure assessments. In the same animals, one wound tissue was used for cellular and molecular studies (n=5), and the other wound for tissue protein and molecular analysis (n=3). We have performed full-excisional wound experiments and functional wound healing analysis on three independent cohorts of mice and each cohort contained 8 animals per group.

### Wound healing assay

Male C57BL/6 (10-week, 20-25 g body weight) mice were purchased from the Jackson Laboratory (Bar Harbor, ME USA). Mice were anesthetized by intraperitoneal injection of ketamine (100 mg/kg) and xylazine (10 mg/kg) body weight, and the hair was removed by shaving and hair removal cream. Skin was disinfected using 70% ethanol and betadine before wounding. The shaved dorsal skin was folded and raised cranially and caudally at midline to generate two, symmetrical, 5-mm diameter wounds using a sterile punch biopsy tool (Robbins instrument, #RBP50) (1, 36, 40). A 5-mm-diameter silicone ring (Grac3 Bio-labs, #CWS-S-0.5) was sutured (DemeTECH, #NYLON5-0) around each wound to restrict contraction. After wound creation and suturing of the silicone ring, wound sites were photographed, and the wound surface was covered with Tegaderm (3M) sterile transparent dressing. After surgery, mice were given SR Buprenorphine (0.05 mg/kg) as post-operative analgesia. Into based on the assigned treatment groups, mice received either saline or 4-AP: saline group (vehicle control), which received 100 μl of saline, and a 4-AP group, which received 40 μg/mouse/daily 4-AP (1.6 mg/kg) in saline intraperitoneally (IP) until day 14 post-wounding. This dosage corresponds with ∼40% of the mouse body surface area and is equivalent of the dosage of 20 mg/day used in treating multiple sclerosis (33, 37, 38, 121) but is less than the dose that have been examined in patients with chronic spinal cord injury. Gross wound healing was monitored daily and images were captured on day 0 to day 14 (days 0, 3, 5, 7, 9, 12 and 14) post-surgery (Figure 1A). Wound areas were measured in pixels using ImageJ-1.53e software (National Institutes of Health, USA) and normalized/corrected for each wound area with reference scales. Wound healing is expressed as percentage with respect to day 0 wounds (1, 36, 40), using the following formula.

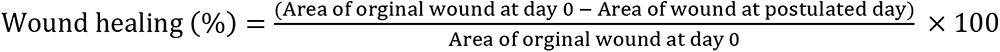

### Histomorphometry analysis

Wound assessments were conducted at 14 days post-wounding using formalin-fixed (Sigma-Aldrich, # HT5011-CS) and paraffin-embedded tissue, serially cut into 5-μm sections on a Microtome (Leica RM2235, Germany). Skin sections were processed for morphometric analysis using hematoxylin and eosin (H&E) (Sigma-Aldrich, #MHS32-1L) and immunofluorescence staining (e.g., (1, 36, 40)). To evaluate the epidermal thickness, and wound-induced hair neogenesis (WIHN), four fields of view per sample were imaged by light microscopy (Olympus BX53, Olympus, Tokyo, Japan) at 10 and 40x magnification. Data were averaged for each mouse and then compared between 4-AP and saline treated groups. Collagen formation, maturation and deposition was carried out using Masson’s trichrome stain (as in, e.g., (122)) as per manufacturer instructions (Sigma-Aldrich, # HT15-1KT). The Masson’s trichome stained slides imaged by light microscopy (Olympus BX53, Olympus, Tokyo, Japan) at 20x magnification and collagen deposition analysis performed using ImageJ-1.53e software.

### Immunofluorescence staining of tissue

Briefly, immunofluorescence analysis was performed on 5-µm thick healed wound sections. The following primary antibodies were used: S100 antibody (Invitrogen, # MA5-12969; 1:200), rabbit nerve growth factor receptor — p75-NTR antibody (MilliporeSigma, # AB1554; 1:500), Chicken neurofilament Heavy-NFH antibody (Novus Biologicals, # NB300-217; 1:500), mouse alpha-smooth muscle actin –α-SMA antibody (Invitrogen, # 14-9760-82; 1:200), mouse TGF-β antibody (Abcam, # 229856; 1:100), rabbit Ki67 antibody (Cell-Signaling, # 9129S; IF-1:400), chicken keratin 15 antibody (BioLegend, # 833901; 1:500), rabbit K17 antibody (gift from Pierre Coulombe to AMN; 1:1000), mouse Cytokeratin14 antibody (Novus Biologicals, # NBP2-34270; 1:100), rat CD31 antibody (BD Biosciences, # 553370; 1:100), mouse anti-SOX10 antibody (Santa Cruz Biotechnology, # sc-365692; 1:100), rabbit vimentin antibody (Thermo Fisher, # 10366-1-AP; 1:200), rabbit NGF antibody (Invitrogen, # MA5-32067; 1:100), keratin-10 (Sigma, # SAB4501656; 1:100), mouse PGP 9.5 antibody (Invitrogen, # PA5-29012; 1:200), and rat substance P antibody (Novus Biologicals, # NB100-65219; 1:100). Sections were incubated overnight with 5% BSA in 0.1% PBS-T for overnight at 4°C. Then incubated with secondary antibodies (Goat anti-Mouse IgG (H+L) or/ Goat anti-Rabbit IgG (H+L) or/Goat anti-Chicken IgG (H+L) or/ Goat anti-rat IgG (H+L) Highly Cross-Adsorbed Secondary Antibody, Alexa Fluor 594 or/488 or/647, #A11008 or/A11032 or/A21449 or/A21247; 1:500) for 1 hour at room temperature. The ProLong™ Gold Anti-fade Mountant with DAPI (Invitrogen, # P36935) was used as nuclear counterstain. The immunofluorescence stained sections were imaged using ZEISS Axio Observer 7-Axiocam 506 mono – Apotome.2 microscope. The image analysis and quantification were carried out using ZEN 2.6 pro (Zeiss) imaging software or ImageJ-1.53e software (National Institutes of Health, USA).

### Human primary cell culture experiments

Foreskin collection and preparation: The human foreskin was rinsed gently with 1X-PBS (ScienCell Research, #0303) containing antibiotic. The hypodermis and blood vessels were removed. Subsequently, the skin was cut into 1-2 mm pieces and then placed in DMEM medium (ScienCell Research, #09221) with dispase-I (Sigma-Aldrich, # D46693) at 4°C for 12-18 hours. After dispase-I treatment, the epidermis was separated from the dermis (66, 67, 123).

Keratinocyte isolation, culture conditions and characterization: The isolated epidermis was placed in a petri dish containing HBSS buffer (Lonza, # CC-5022) for 10 minutes at room temperature, then treated with trypsin (Lonza, # CC-5012) at 37°C until the epidermis became loose and the medium cloudy due to keratinocyte release. The cloudy medium was collected and trypsin activity neutralized using fetal bovine serum (FBS, ThermoFisher Scientific # 10082147) in 1:1 ratio. The epidermis and suspended keratinocytes were centrifuged at 1500 rpm for 5 minutes. The pellet was resuspended in KGM-GOLD keratinocyte medium (Lonza KGM gold and supplements, # 00192151 and 00192152) (65, 67, 123). The isolated foreskin was cultured at 37°C in a 5% CO_2_ incubator for 1-2 days to allow keratinocytes to adhere. Adherent keratinocytes were maintained in KGM-Gold medium until cells reached about 80% confluent.

Dermal Schwann cell isolation, culture conditions and characterization: The separated dermis was minced into small pieces and placed in a petri dish containing collagenase (Gibco, # 17018-029) in DMEM basal medium at 37°C for 2.5 hours. The dermis was then dissociated cells were collected and centrifuged at 1500 rpm for 5 minutes. The pellet was resuspended in complete DMEM medium, and cells were plated on poly-L-lysine (ScienCell Research, #0403) coated dished at 37°C in a 5% CO_2_ incubator overnight. The next day, adherent cells were treated with 10 µM cytosine arabinoside (Sigma-Aldrich, # C1768) containing DMEM complete medium and incubated at 37°C in 5% CO_2_ incubator for 24 hours. After this treatment, the cells were cultured in Schwann cell culture medium (ScienCell Research, #1701) (66, 123) until cells reached ∼95% confluency.

### Immunofluorescence staining of cells

Indirect immunofluorescence analysis was used to identify, characterize and analyze proliferation after 4-AP treatment. For each cell type (keratinocytes, and Schwann cells) treated and untreated cultures were processed in the same experimental session. An equal number (1 × 10^4^ cells/chamber) of passage 1 cells were seeded on chamber slides (Lab-Tek, #154526). The cells were grown in their respective complete medium in the presence or absence of 4-AP for 72 hours. Cells then were fixed with 4% paraformaldehyde (ThermoFisher, # J19943-K2) followed by 0.1% tritonX-100 and stained with primary antibodies used against, mouse Cytokeratin14 antibody (Novus Biologicals, # NBP2-34270; 1:200), keratin-10 (Sigma, # SAB4501656; 1:100), S100 antibody (Invitrogen, # MA5-12969; 1:200), rabbit nerve growth factor receptor — p75-NTR antibody (MilliporeSigma, # AB1554; 1:500) in 5% BSA-containing PBS (as in, e.g., (65–67, 123)). After washing, cells were incubated with respective secondary antibodies for 1 hour at room temperature. The ProLong™ Gold Anti-fade Mountant with DAPI (Invitrogen, # P36935) was used as nuclear counterstain. The immunofluorescence stained sections were imaged using ZEISS Axio Observer 7-Axiocam 506 mono – Apotome.2 microscope.

### Cell viability assay with 4-aminopyridine

The keratinocytes, and Schwann cells were cultured in 96-well plates for 18 hours. Cells were placed in minimal media (no serum or growth factors) for 4 hours prior to 4-AP treatment. Cells were treated with 4-AP (at concentrations ranging from 1 to 10000 µM) in appropriate cell culture medium for 24 hours. The cell viability following 4-AP treatment was assayed by MTT assay according to manufacturer’s protocols (Roche, cell proliferation kit I (MTT), # 11465007001). The percentage of live keratinocytes, Schwann cells was determined using the following formula.

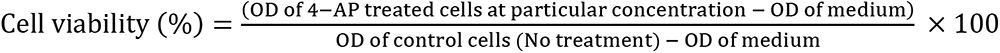

### Cells scratch wound healing migration assay

Keratinocytes, and Schwann cells (7×10^4^ cells/well keratinocytes and 3.5×10^4^ cells/well of SCs), in their respective media, were seeded on tissue culture dishes pre-coated either with collagen-I (corning life science, # 354236) or PLL on 96-well ImageLock microplates for 6 hours (Incucyte-sartorius plate, # 4379). For drug treatment, cells were pretreated with 4-AP for 18 hours prior to perform the wound scratch assay. Next, wound scratches were created using the IncuCyte automated system (Essen BioScience) (e.g., (59, 69)). After scratching, the cells were washed with PBS and the respective cell media was added with or without 1 mM of 4-AP. The plate was incubated in the IncuCyte™ automated imaging system, and wound healing and cell migration was monitored by time-lapse photography capturing images every hour from 0 to 42 hours. The relative area of wound size and cell migration at each time point was analyzed using the IncuCyte™ Scratch Wound Cell Migration Software Module (Essen BioScience) and the percent of wound healing was calculated from the area measured after scratching relative to the basal area as expressed in pixels.

### Co-culture of keratinocytes with Schwann cells in wound scratch assays

To determine the influence of Schwann cells on keratinocyte migration and proliferation wound healing scratch assays were conducted. The co-culture experiment was performed with keratinocytes combined with either Schwann cells in a ratio of 10:1. The same propositions of their respective media were added and cells were seeded on 96-well ImageLock microplates for 6 hours, in wells pre-coated with collagen-I. For the drug treatment, cells were pretreated with 4-AP prior to plating on collagen-coated for 18 hours. Next, wound scratches were created and cells were grown with or without 4-AP in the medium. Cells were imaged every hour until 24 hours and analyzed for percent of closure as described above.

### Tissue protein isolation and western blot analysis

For protein isolation the harvested skin tissue was flash frozen immediately. The frozen skin tissue was ground to a fine power using a liquid nitrogen mortar. The harvested cells and /or tissue powder was dissolved in RIPA buffer containing Halt™ Phosphatase (Thermo Scientific, # 78420) and Protease Inhibitor Cocktail (Roche complete tablets mini EASYpack, # 04694124001). Tissue and cell debris were removed by centrifugation at 14000 rpm for 30 minutes at 4°C. The supernatants were collected and the total protein concentration was determined by BCA protein assays (Thermo Scientific™ Pierce™, # 23225). The proteins (20-30 μg) of the tissue protein samples were subjected to 12% sodium dodecyl sulfate polyacrylamide gel electrophoresis (Bio-Rad mini-PROTEAN TGX Gels, # 4561044) and transferred to polyvinylidene fluoride (PVDF) membranes. After the membranes were blocked with 5% non-fat milk in 1X TBS-T for 1 hour, they were incubated with the appropriate primary antibodies (rabbit nerve growth factor receptor — p75-NTR antibody (MilliporeSigma, # AB1554; 1:500), mouse alpha-smooth muscle actin –α-SMA antibody (Invitrogen, # 14-9760-82; 1:500), rabbit TGF-β antibody (Cell Signaling Technology, #3711S, 1:500), mouse anti-SOX10 antibody (Santa Cruz Biotechnology, # sc-365692; 1:100), rabbit NGF antibody (Invitrogen, # MA5-32067; 1:1000), Mouse GAPDH antibody (ThermoFischer, #MA5-15738; 1:1000) at 4°C overnight, then incubated with HRP-conjugated secondary antibodies (dilution,1:3000) for 1 hour. Immunoreactivity was then detected using chemiluminescent substrate (Thermo Scientific™ SuperSignal™ West Pico PLUS, # 34577). The intensities of the bands were quantified using Gel-imaging software (Bio-Rad Laboratories Inc., Image Lab 6.1). The quantified band intensities were normalized using GAPDH and expressed either as normalized intensity or as ratios with respect to saline treated mice.

### Statistical analysis

The number of animals per group were determined by pre-hoc power analysis of preliminary data by a qualified statistician to achieve at least 80% power for the primary outcome in this study. Animals were randomly assigned into either saline treatment groups. All results are presented as mean ± standard error of the mean (SEM). Data were analyzed either by two-tailed Sidak’s for wound healing functional analysis multiple time-point comparison and unpaired data from different experiments by one-way ANOVA followed by unpaired *t* test and nonparametric test after confirmation of normally distributed data. Statistical analysis was performed using the GraphPad PRISM 9.2.0(332) (GraphPad, La Jolla, CA, USA) and star indicates statistical significance of * = *P* between 0.01 and 0.05, ** = *P* between 0.01 and 0.001, *** = *P* between 0.001 and 0.0002, and **** = *P* between 0.0002 and 0.0001 versus saline considered as significant.

### Study approval

Mouse studies were carried out in accordance with the NIH’s Guide for the Care and Use of Laboratory Animals (NIH publication No. 86–23, revised in 2011) and the animal protocol was approved by The Penn State College of Medicine Institutional Animal Care and Use Committee (IACUC No.: PROTO202001314). All the animal experimental procedures were performed in accordance with the Penn State College of Medicine institutional guidelines for animal care established by The Pennsylvania State University.

## Acknowledgments

This work was supported by grants from the NIH (K08 AR060164-01A) and DOD (W81XWH-16-1-0725) to JCE. in addition to institutional support from The Pennsylvania State University Hershey Medical Center. We thank Dr. Hong-Gang Wang and Ms. Xiaoming Liu for allowing us to use the IncuCyte™ instrument and technical assistance. We would like to thank Dr. Fadia Kamal, Dr. Reyad Elbarbary, and Dr. Srinivas Koduru for allowing us to use lab instruments and technical support. We thank Dr. Pierre A Coulombe for providing the K17 antibody. We would also like to acknowledge staff technical assistance from the Penn State College of Medicine Animal Facility and the Center for Orthopaedic Research and Translational Science (CORTS).

## Author contributions

JCE conceived the study. MGJ, PKG, AMN, and JCE designed the study. MGJ, PKG, and JCE performed experiments and were involved in the acquisition, data analysis. MGJ, PKG, AMN, MDN, and JCE involved in the interpretation of data. AMN, and JCE contributed reagents/materials/analysis tools. MGJ, AMN, MDN, and JCE wrote the manuscript. All the authors read and edited the manuscript and have given approval to the final version of the manuscript.

## Notes

### Competing Interest Statement

MGJ, PKG, JCE- are inventors on patent/patent application (63/005,890; 63/133,407; PCT/US21/14442; and PCT/US22/11351) submitted by Pennsylvania State University that covers Method and materials for treating nerve injury and/or wound healing and Therapeutic treatments for hair growth. JCE and MN are founding members of Solaxa Therapeutics, a company that has licensed a previous patent on the use of 4-AP for traumatic peripheral nerve injury and nerve continuity diagnosis. All other authors declare that they have no competing financial interests.

