## Supplemental Data for "4-aminopyridine promotes accelerated skin wound healing via neurogenic-mediator expression"

**Supplemental Figures**

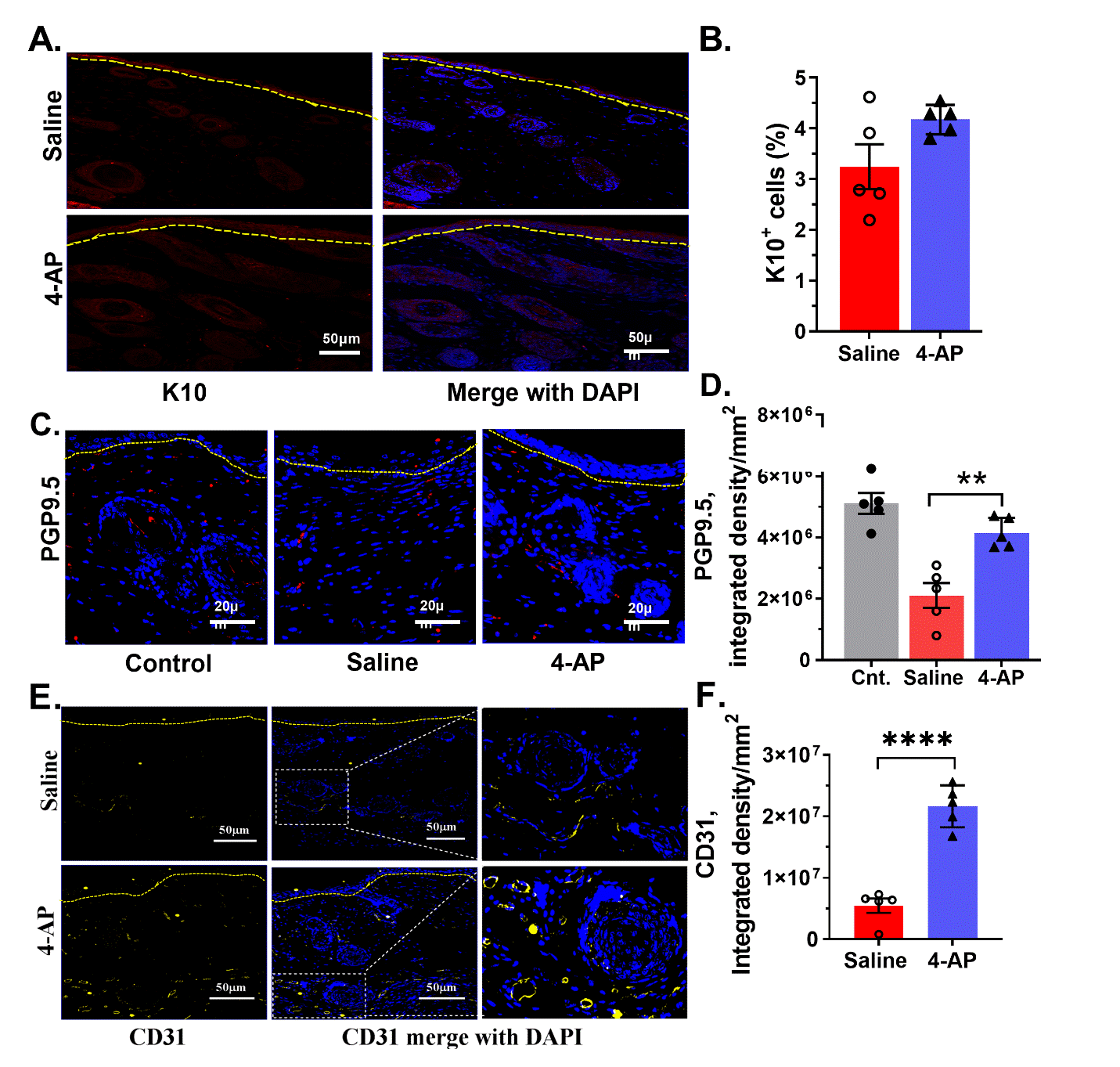

**Supplemental Figure 1. 4-AP induced neo-angiogenesis and neuronal peptide wound healing and did not alter keratinocyte K10 expression. (A)** Keratin 10 protein expression in healed epidermis by immunofluorescence. 4-AP treatment did not cause any change in expression of keratinocyte K10 expression. K10 (red); DAPI (blue) denotes nucleus and dashed line denotes epidermal/derma border. Scale bars, 50 µm. **(B)** Percent of K10^+^ cells in control and 4-AP treated skin wounds at day 14. Each image represents 20 images from 5 different mice wound tissue and data represented as mean ± SEM, *n=5* animals per group. **(C)** Immunofluorescence staining of control and healed wound sections for pan-neuronal marker PGP-9.5 (red) and nuclear stain DAPI (blue) denotes nucleus and dashed line denotes epidermal/derma border. Scale bars, 20 μm. **(D)** Quantification of PGP-9.5 protein expressing cells showed significantly increased PGP-9.5 intensity in the 4-AP treated group compared to the saline treated group at day 14. PGP 9.5 in 4-AP-treated mice was not significantly different from seen in uninjured (control) tissue. Each image represents 20 images from 5 different mouse wounds and data are represented as mean ± SEM, *n = 5* animals per group, statistical significance indicated by asterisks (** = P between 0.01 and 0.001 vs. saline). **(E)** CD31 protein expression in healed skin by immunofluorescence. CD31 (yellow); DAPI (blue) denotes nucleus and dashed line denotes epidermal/derma border. Scale bars, 50 µm. **(F)** Quantification of CD31 staining intensity, 4-AP treated group showed higher intensity compared to saline control group. Each image represents 20 images from 5 different mouse wounds and data are represented as mean ± SEM, *n = 5* animals per group, statistical significance indicated by asterisks (**** = P between 0.0002 and 0.0001 vs. saline).

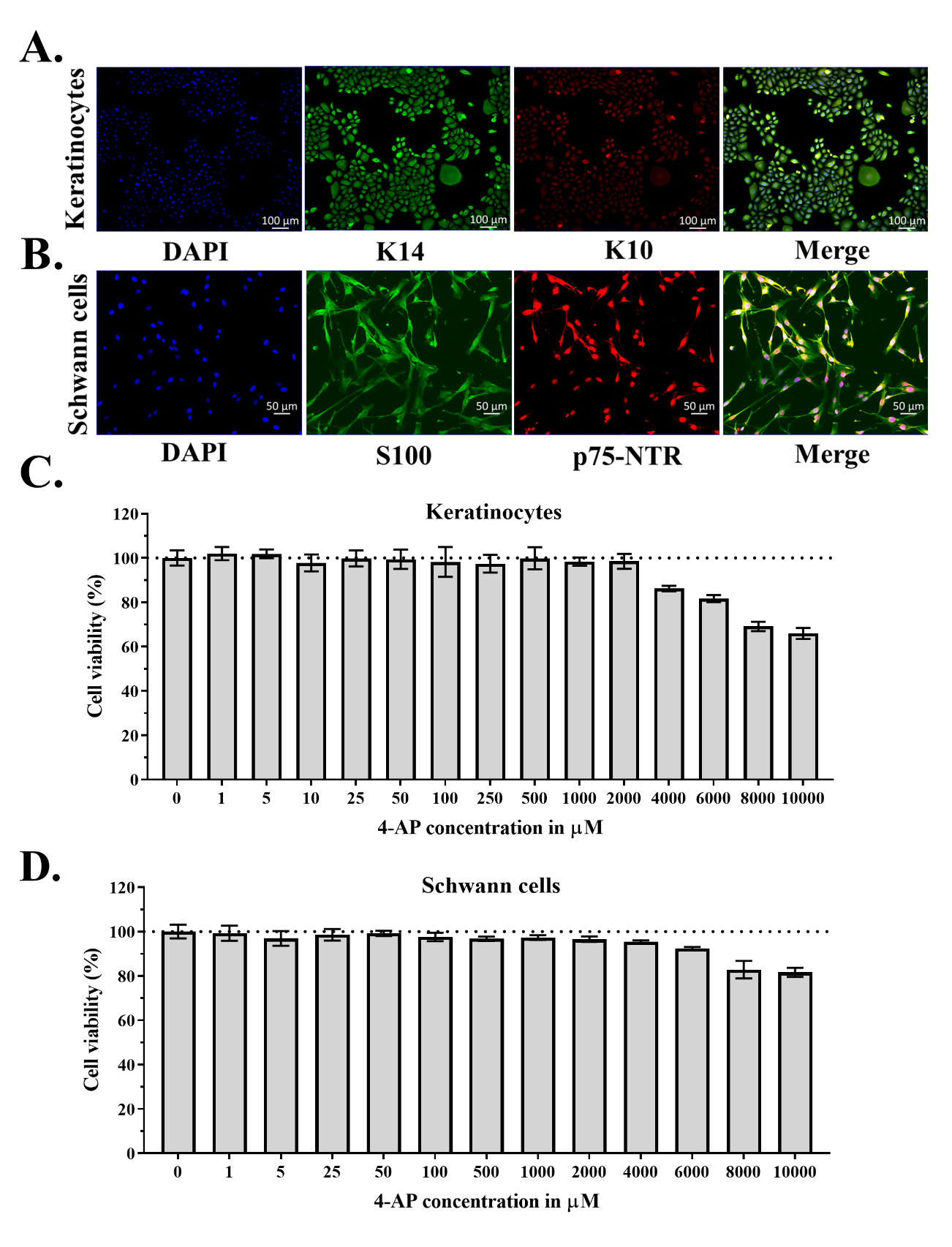

**Supplemental Figure 2. Isolation and characterization of human skin derived primary keratinocytes, and Schwann cells.** (A) Keratinocytes were characterized using keratin 14 (K14 (green), a marker of proliferative keratinocytes), K10 (keratin 10 (red), a marker of keratinocyte differentiation marker) and DAPI (blue). Scale bars, 100µm. **(B)** Schwann cells were identified using S100 (green, Schwann cells marker), p75-NTR (red, nerve growth factor receptor marker) and DAPI (blue). Scale bars, 50 µm. **(C-D)** Cell viability using MTT assay with different concentrations of 4-AP (ranging from 1 to 10000 µM) for **(C)** Keratinocytes, **(D)** Schwann cells and data represented as mean ± SEM, *n=3* replicates/concentration of 4-AP.

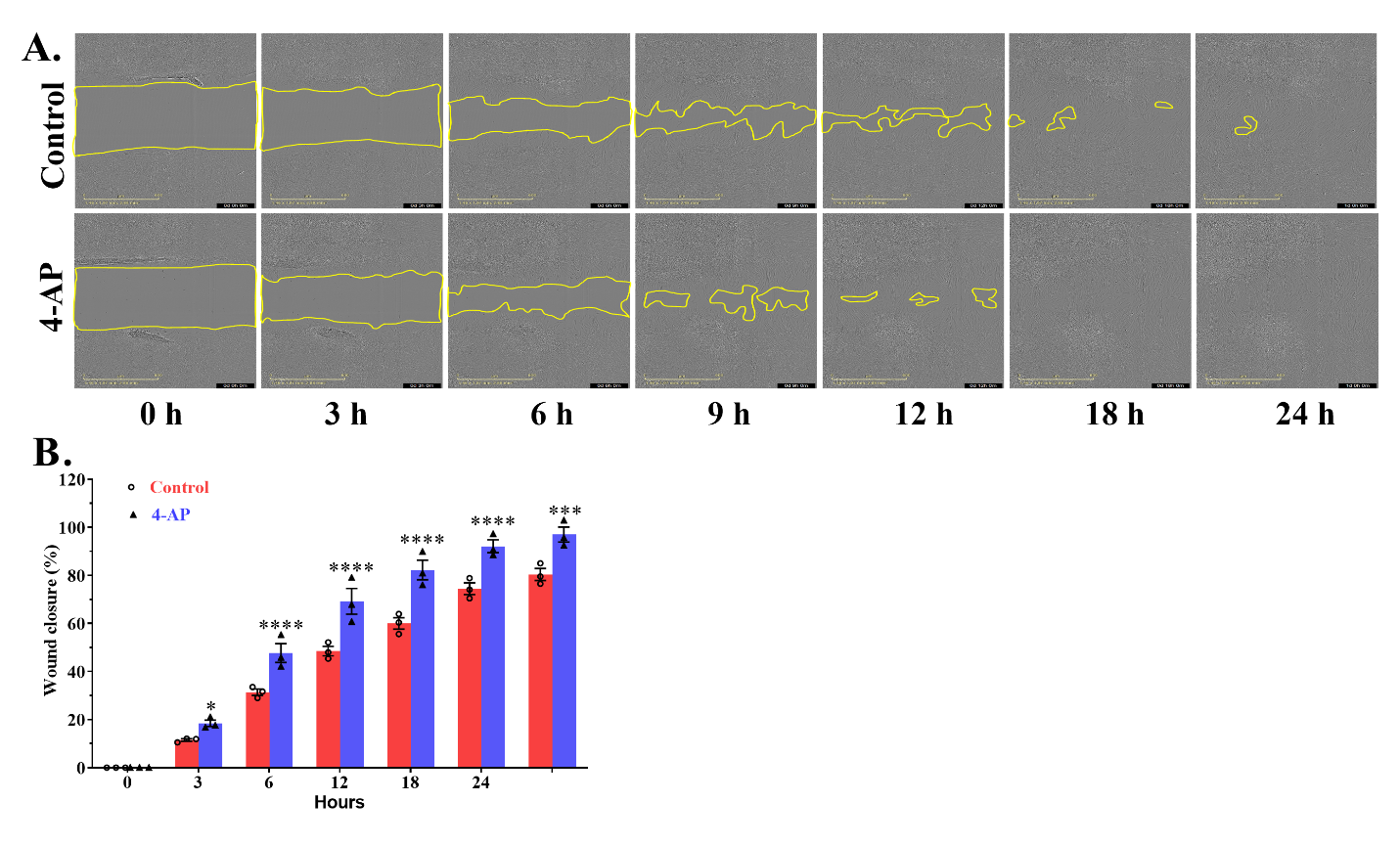

**Supplemental Figure 3. Effect of 4-AP exposure on primary dermal Schwann cell in scratch wound healing assays.** **(A)** Representative images of *in-vitro* Schwann cells scratch assays with 4-AP and vehicle controls at indicated time points. Scale bar, 100 µm. **(B)** The relative percentage of wound closure was calculated as the ratio of the remaining wound gap at the given time point compared to time 0. Each image represents 9 images from 3 biological replicates and data represented as mean ± SEM, *n=3* biological replicates per group, statistical significance indicated by asterisks (* = P between 0.01 and 0.05, *** = P between 0.001 and 0.0002, and **** = P between 0.0002 and 0.0001 vs. control).

**Supplementary Movies**

**Supplemental Movie 1.** An example time-lapse phase contrast images depicting the migration video of keratinocytes without treatment (control) during wound scratch closure. Images were recorded every one hour. Scale bar, 100 µm.

**Supplemental Movie 2.** An example time-lapse phase contrast images depicting the migration video of keratinocytes after 4-AP treatment during wound scratch closure. Images were recorded every one hour. Scale bar, 100 µm.

**Supplemental Movie 3.** An example time-lapse phase contrast images depicting the migration video of Schwann cells without treatment (control) during wound scratch closure. Images were recorded every one hour. Scale bar, 100 µm.

**Supplemental Movie 4.** An example time-lapse phase contrast images depicting the migration video of Schwann cells after 4-AP treatment during wound scratch closure. Images were recorded every one hour. Scale bar, 100 µm.

**Supplemental Movie 5.** An example time-lapse phase contrast images depicting the migration video of co-cultured keratinocytes and Schwann cells without treatment (control) during wound scratch closure. Images were recorded every one hour. Scale bar, 100 µm.

**Supplemental Movie 6.** An example time-lapse phase contrast images depicting the migration video of co-cultured keratinocytes and Schwann cells after 4-AP treatment during wound scratch closure. Images were recorded every one hour. Scale bar, 100 µm.
